# α-Synuclein Aggregation is Triggered by Amyloid-β Oligomers via Heterogeneous Primary Nucleation

**DOI:** 10.1101/2022.06.20.496547

**Authors:** Devkee M. Vadukul, Marcell Papp, Rebecca J. Thrush, Jielei Wang, Yiyun Jin, Paolo Arosio, Francesco A. Aprile

## Abstract

An increasing number of cases where amyloids of different proteins are found in the same patient are being reported. This observation complicates diagnosis and clinical intervention. Amyloids of the amyloid-β peptide or the protein α-synuclein are traditionally considered hallmarks of Alzheimer’s and Parkinson’s diseases, respectively. However, the co-occurrence of amyloids of these proteins has also been reported in patients diagnosed with either disease. Here, we show that soluble species containing amyloid-β can induce the aggregation of α-synuclein. Fibrils formed under these conditions are solely composed of α-synuclein to which amyloid-β can be found associated, but not as part of the core of the fibrils. Importantly, by global kinetic analysis, we found that the aggregation of α-synuclein under these conditions occurs via heterogeneous primary nucleation, triggered by soluble aggregates containing amyloid-β.

## INTRODUCTION

Neurodegenerative diseases, such as Alzheimer’s (AD) and Parkinson’s (PD) diseases, are characterized by the formation of fibrillar protein aggregates, called amyloids, in the nervous system (1). Amyloids are enriched in cross-β sheet and resistant to degradation (2,3). In the case of AD, a key pathological hallmark are the extracellular amyloid plaques made of the amyloid-β peptide (Aβ) (4). Aβ exist as variants of different lengths (i.e., 38-49 residue-long) and originate by the proteolytic cleavage of the amyloid protein precursor (5-7). Aβ monomers undergo a self-assembly process that leads to the formation of amyloids via small transient intermediates, called oligomers (4). Thus far, Aβ monomers and amyloids are considered less pathogenic compared to the oligomers (8-11). Several toxic effects of Aβ oligomers have been identified, including inflammation, synaptotoxicity, membrane permeabilization and oxidative stress (8,9,12-19).

The co-occurrence of amyloids composed of Aβ and of other proteins has been detected in several neurodegenerative diseases, suggesting an interplay between Aβ and these amyloid forming proteins (20-26). One such protein is α-synuclein (α-syn), which is found as amyloid fibrils in Lewy bodies (LBs) in PD, Lewy body dementia, and Multiple System Atrophy (27-29). The identification of Aβ plaques in up to 50% of PD patients (30,31) and of LBs in almost 50% of AD patients (32) suggests a crosstalk between Aβ and α-syn. This hypothesis is further supported by the identification of the non-amyloid-β component (NAC) region of α-syn in AD plaques (32,33). Additionally, as Aβ has been found to accumulate intracellularly (34), where α-synuclein is found (35), it is likely that the two proteins interact within the cellular environment.

The effects of Aβ and α-syn on each other’s aggregations are not yet fully characterized at the molecular level. Recently, it has been found that α-syn can either accelerate or inhibit the aggregation of Aβ. This dual mechanism of α-syn has been observed on both the 40-residue-long (Aβ40) and the 42-residue-long (Aβ42) isoforms of Aβ (22,36-40). Whether α-syn promotes or inhibits Aβ aggregation seems to depend on the aggregated state of α-syn. In fact, it has been shown that α-syn monomers can delay the aggregation of Aβ42, whilst α-syn amyloid fibrils can accelerate it (41). Conversely, it has been shown that Aβ42 is able to trigger the aggregation of α-syn (42). Furthermore, the formation of hetero-oligomers made of both Aβ42 and α-syn has been observed *in vitro*, suggesting that the two proteins can co-aggregate and form heterogeneous aggregates (43-45). However, to date, there is only limited information on the mechanism by which Aβ induces the aggregation of α-syn or the structure of the aggregates which are formed when the two proteins are present together.

Here, we investigate the structure and the kinetics of formation of aggregates formed when α-syn and Aβ42 co-aggregate *in vitro*. We report that these aggregates are α-syn fibrils, whose surface is coated by Aβ42. The mechanism of formation of these fibrils can be described with a model where Aβ42 oligomers provides α-syn monomers with a surface to nucleate, similarly to what has been seen with lipids (46). Our results provide a first mechanistic understanding on how these two key amyloidogenic proteins can possibly interact when found co-aggregated in patients. They also reveal a new toxic mechanism of Aβ oligomers, namely their ability to trigger the formation of amyloids of other proteins.

## RESULTS

### Fibrils formed in α-syn–Aβ42 co-incubation are made of α-syn

To assess the effects of Aβ42 on α-syn aggregation, we carried out thioflavin T (ThT) assays on solutions containing 60 μM monomeric α-syn in the presence of a sub-stoichiometric concentration (2 μM) of monomeric Aβ42 (**Fig. 1a and Fig S1a-c**). Hereafter, we will refer to this condition as the *co-incubation sample*. As controls, we also performed ThT experiments on solutions containing either 60 μM α-syn or 2 μM Aβ42. We found that α-syn alone did not aggregate during the time of this experiment (8 h), while Aβ42 alone aggregated in a timescale of approximately 4 h as previously reported (47). In the co-incubation sample, we observed an increase of the ThT signal, which was slower than the aggregation profile of Aβ42 alone. In fact, the aggregation half-time (t_50_) of the co-incubation sample was approximately 1.5-fold the one of Aβ42 alone (3 ± 0.4 and 2 ± 0.4 h, respectively) (**Fig. S1d)**. Furthermore, by negative stain TEM, we confirmed that the aggregates formed in the co-incubation had a fibrillar conformation (**Fig S1d**).

**Figure 1.**
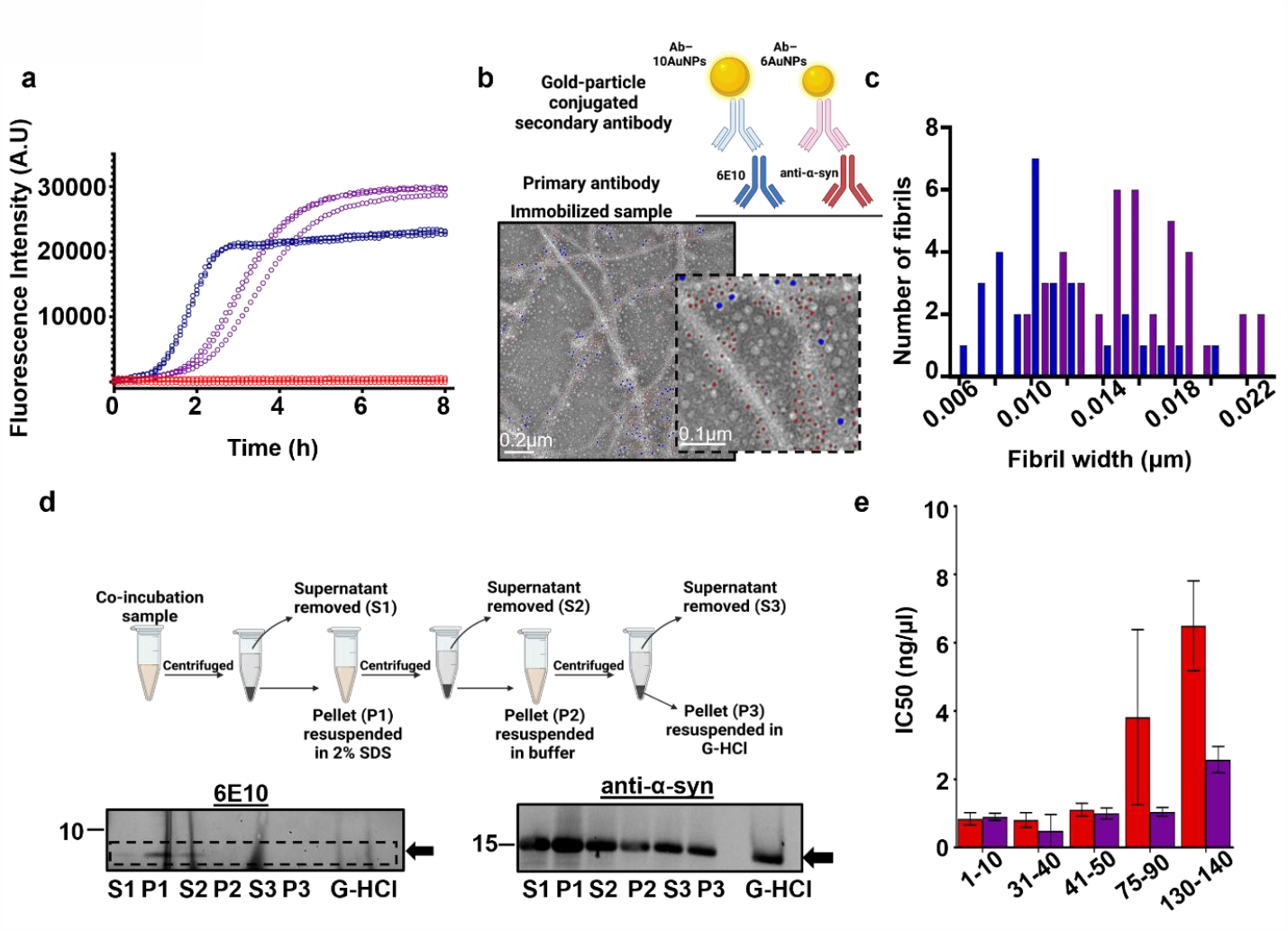
Aggregation of α-syn–Aβ42 co-incubation results in α-syn fibrils. (a) ThT fluorescence assay of 60 μM α-syn (red), 2 μM Aβ42 (blue) and α-syn aggregated with Aβ42 (purple). Three replicates are shown per condition. (b) Immunogold labelling and negative stain electron micrograph α-syn aggregated with Aβ42 at the end point of aggregation. Fibrils are highly decorated with 6nm gold particles (Ab–6AuNPs, red arrows) specific for α-syn as opposed to 10nm gold particles labelling Aβ (Ab–10AuNPs, blue arrows) which are sparse. (c) Width distribution of Aβ42 fibrils (blue, n = 32) and fibrils in the co-incubation sample (purple, n = 32). (d) Schematic of G-HCl treatment of SDS insoluble fraction (top). Fibrils from the co-incubated sample were collected by centrifugation and the supernatant (S1) was removed and separated from the pellet (P1). The pellet was washed with 2% SDS and centrifuged. The supernatant was again removed (S2) and the pellet was resuspended in buffer (P2). The pellet was centrifuged, and the supernatant (S3) and pellet were separated (P3) after which the pellet was treated with 4M G-HCl for 2h at RT before SDS-PAGE and western blot analysis. Detection with the 6E10 antibody (bottom left) revealed no Aβ42 in the G-HCl treated pellet, however, α-syn was detected with the anti-α-syn antibody (bottom right). Monomers of the proteins are indicated by the black arrows (e) Proteinase K digestion of α-syn fibrils formed in the absence (red) and presence (purple) of Aβ42 quantified by densitometry and SDS-PAGE and western blotting. Quantification from two independent experiments. Error bars representative of SEM.

To investigate the protein composition of the fibrils from the co-incubation sample, we performed immunogold TEM on samples collected at the aggregation endpoint (24 h) (**Fig. 1b** and **Fig. S2**). To do so, the samples were simultaneously probed for Aβ42 with antibodies conjugated to 10 nm gold nanoparticles (Ab–10AuNPs, blue) and α-syn with antibodies conjugated to 6 nm gold nanoparticles (Ab–6AuNPs, red). We found no or extremely low signal for Aβ42 in the sample with α-syn alone and for α-syn in the samples with Aβ42 alone, verifying that our staining is specific (**Fig. S2**). The sample containing α-syn alone showed a diffuse distribution of small clusters of Ab–6AuNPs, confirming the presence of soluble α-syn and the lack of fibrils (**Fig. S2c**), similarly to previous reports (42) and in agreement with our ThT and negative staining TEM data. The sample with Aβ42 only contained fibrils decorated by Ab– 10AuNPs, i.e., made of Aβ42 (**Fig. S2a**). Fibrils were also present in the co-incubation sample (**Fig. 1b**). However, these fibrils were structurally distinct from Aβ42 fibrils. The median of the widths of the fibrils from co-incubation (16 nm) was larger than the one of Aβ42 fibrils (10 nm) (**Fig. 1c**). Furthermore, the fibrils in the co-incubation sample were predominantly decorated by 6Ab–AuNPs, i.e., made of α-syn. Some Ab–10AuNPs could also be detected on these fibrils. To understand whether Aβ42 was peripheral or within the SDS-insoluble core of the fibrils, we quantified Aβ42 and α-syn in the SDS-insoluble protein fraction at the aggregation endpoint. To do so, we washed the fibrils with SDS to remove any associated soluble aggregates and then treated the fibrils with 4M guanidine hydrochloric acid (G-HCl). This was then analyzed by SDS-PAGE and western blotting (**Fig. 1d**). We found that only α-syn was in the SDS-insoluble fractions, indicating that the core of the fibrils from the co-incubation sample is composed of α-syn only, and Aβ42 is on the surface but not within the core of the fibrils.

To further investigate the stability of the co-incubation fibrils, we treated them with varying concentrations of proteinase K (PK) and measure the PK-resistant α-syn by SDS-PAGE and western blotting as well as densitometry using an array of antibodies scanning the sequence of α-syn (**Table S1**). We found that the PK-stability profile of the fibrils in the co-incubation sample was largely comparable to the one of fibrils obtained by incubating α-syn for two weeks. However, based on the profile of degraded protein below the monomer band, the fibrils formed from α-syn alone have a more resistant NAC region (probed with the antibody recognizing amino acids 75-90), which makes up the core of α-syn fibrils (**Fig. 1e and S3**). This analysis further confirms the amyloid nature of the fibrils in the co-incubation sample and indicates α-syn fibrils formed in the presence of Aβ42 are structurally distinct to α-syn fibrils formed in the absence of Aβ42.

### α-syn aggregation in α-syn–Aβ42 co-incubation is triggered by Aβ42 oligomers

We then aimed to identify the conformation(s) of Aβ42 that trigger α-syn aggregation.

We performed immuno-dot blots on soluble protein fractions at the start and endpoint (24 h) of aggregation (**Fig. 2a and S4**). As expected, we observed that soluble Aβ42 drastically decreases (by ∼70%) when incubated alone. On the contrary, in the co-incubation sample, the soluble protein species does not significantly change. Soluble α-syn decreases by only ∼10% when the protein is incubated alone, while, in the co-incubation sample, it reduces by ∼30%. By native-PAGE and western blotting on protein total fractions (**Fig. 2b**), we found that, during aggregation, Aβ42 alone progressively forms protein species that do not enter the gel, which is a feature of fibrils. Instead, α-syn alone remains mainly monomeric. In the co-incubation sample, more Aβ42 can enter the gel as compared to the Aβ42 alone condition. In particular, the protein forms high molecular weight oligomers after 4 h of incubation. Additionally, higher molecular weight assemblies of α-syn are not detected until Aβ42 oligomers are formed. Together, these data indicate that Aβ42 oligomers are triggering α-syn aggregation.

**Figure 2.**
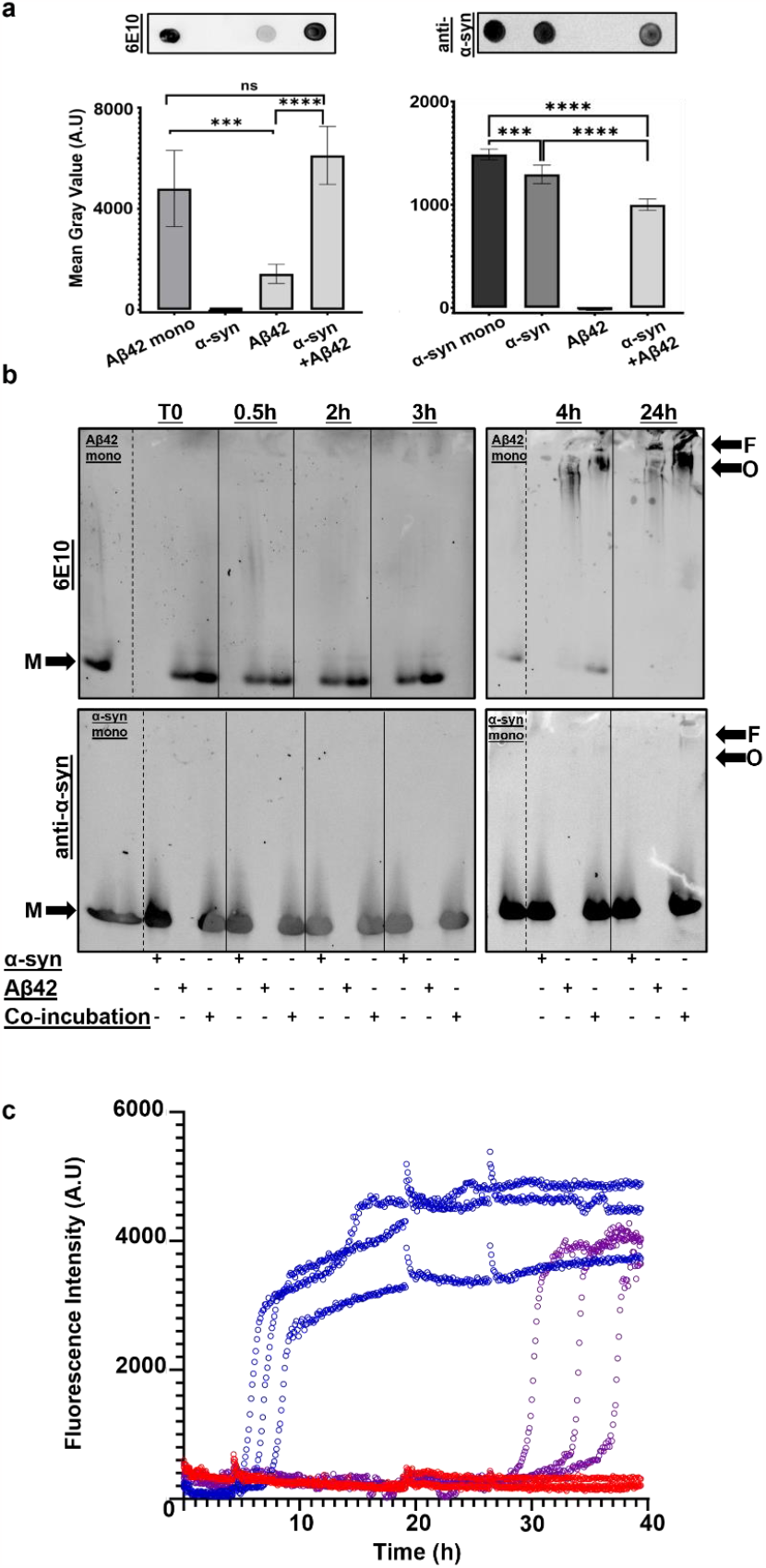
Aβ42 oligomers trigger α-syn aggregation. (a) Dot blot analysis and quantifications on the soluble fractions of aggregated samples detected with 6E10 (left) and anti-α-syn (right) primary antibodies. 5 repeats for each sample were quantified. Error bars are shown as SD. Mean grey values were compared with One-way ANOVA, Tukey’s multiple comparison test where p = 0.1234 (ns), 0.0332 (*), 0.0021 (**), 0.0002 (***) and 0.0001 (****). (b) Native-PAGE and western blot analysis of total samples during aggregation. Detection with the 6E10 antibody (top) and anti-α-syn (bottom). Our results show that higher molecular weight assemblies of α-syn (O and F) are not seen until oligomers of Aβ42 are formed at 4h, as compared to the monomers of both proteins (M). (c) ThT fluorescence assay of 2 μM Aβ42 (blue), 60 μM α-syn (red) and the co-incubation sample (purple) in low phosphate buffer (4mM). Three individual technical replicates are shown.

To understand whether monomeric or aggregated Aβ42 is responsible for triggering α-syn aggregation, we performed ThT experiments under conditions where Aβ42 aggregation is delayed. Our hypothesis was that if Aβ42 aggregated forms, but not the monomers, are responsible for triggering α-syn aggregation, then conditions which delay the aggregation of Aβ42 also delay α-syn aggregation when the two proteins are co-incubated. Thus, we performed ThT experiments in 4mM phosphate buffer instead of 20mM phosphate buffer (**Fig. 2c)**. We found that α-syn does not aggregate in low phosphate conditions similarly to when it is incubated in 20mM phosphate buffer. Aβ42 aggregates significantly slower in 4mM phosphate buffer compared to 20mM phosphate buffer. In co-incubation conditions, α-syn aggregation is also significantly delayed, suggesting that a direct interaction between α-syn and aggregated, but not monomeric, Aβ42 plays a key role in the process.

To rule out the possibility that Aβ42 fibrils can also induce α-syn aggregation, we monitored the aggregation that occurs when Aβ42 fibrils are co-incubated with α-syn (**Fig. 3 and S5)**. We measured the ThT fluorescence of samples containing α-syn alone, Aβ42 fibrils alone, and α-syn in the presence of Aβ42 fibrils (**Fig. 3a**). As expected, we observed that α-syn alone does not aggregate. Aβ42 fibrils alone have a steady ThT fluorescence as does the co-incubation sample. Immunogold TEM showed that the fibrils in the co-incubation sample are largely labelled with the Ab–10AuNPs specific for Aβ (**Fig. 3b and S5a**). This result confirms that there is no aggregation of α-syn and, conversely, that Aβ42 fibrils are not affected by soluble α-syn. Native-PAGE and western blotting on the endpoints of aggregation (**Fig. 3c**) revealed no soluble Aβ42 in the Aβ42 alone and co-incubation samples. No difference in the molecular weight of α-syn was observed in the α-syn alone and co-incubation samples. This evidence indicates that fibrillar Aβ42 does not affect the aggregation of α-syn or vice versa. Strengthening this observation, dot blot analysis on the soluble and insoluble protein fractions revealed that no soluble Aβ42 was present in the sample containing Aβ42 fibrils alone (**Fig. 3d and S5b**). As expected, soluble Aβ42 was not detected in the co-incubation sample confirming that α-syn had no destabilizing effect on fibrillar Aβ42 Additionally, no difference in the amount of soluble α-syn between the co-incubation and α-syn alone samples was detected Based on these data, we rationalize that soluble Aβ42 oligomers but not the monomers or fibrils trigger the aggregation of α-syn.

**Figure 3.**
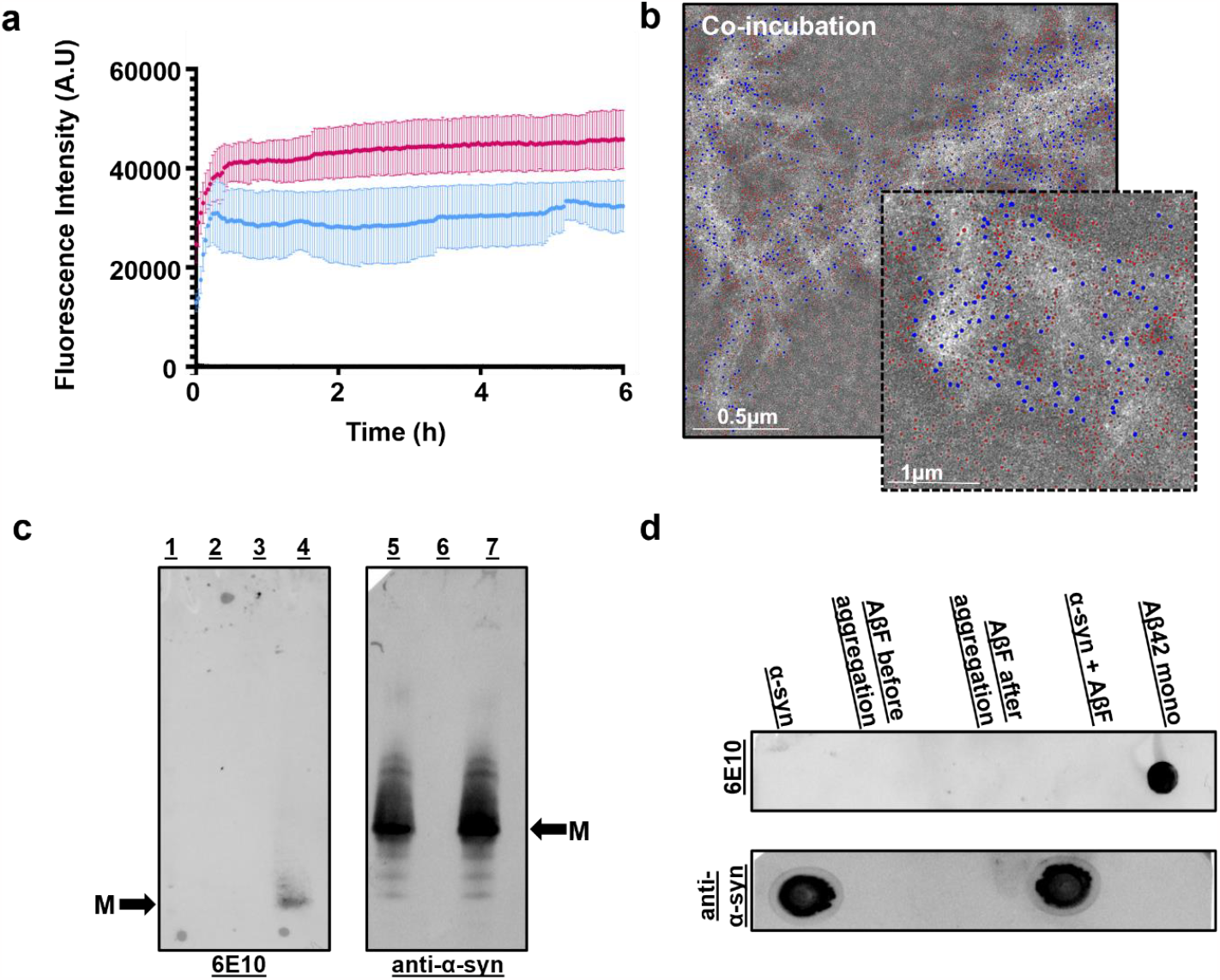
Aβ42 fibrils do not trigger the aggregation of α-syn. (a) ThT fluorescence assay of 60 μM α-syn (black), 2 μM Aβ fibrils (blue) and α-syn aggregated with Aβ fibrils (pink). The average of three replicates for each condition are shown. Error bars represent standard deviation. (b) Immunogold labelling and negative stain TEM of α-syn aggregated with Aβ fibrils and Aβ fibrils alone at the end point of aggregation. Fibrils are highly decorated with 10nm gold particles (Ab–6AuNPs, red) specific for Aβ42 as opposed to 6nm gold particles labelling α-syn (Ab–10AuNPs, blue) which are largely in the surrounding area of the fibrils. (c) Native-PAGE and western blot analysis of total samples after aggregation. Detection with 6E10 (left) revealed no detection of (1) α-syn, (2) that Aβ42 fibrils alone do not enter the gel, (3) the co-incubated sample also has no detectable Aβ42 in the gel however (4) freshly purified Aβ42 monomers are detectable. Detection with the anti-α-syn (right) revealed no difference in the molecular weight assemblies of α-syn either (5) alone or (7) aggregated with Aβ42 fibrils. (6) Aβ42 fibrils are not detected by this antibody. (d) Dot blot analysis on the soluble fractions of aggregated samples detected with 6E10 (left) and anti-α-syn (right) primary antibodies. No soluble Aβ42 was detected in any sample except freshly purified Aβ42 monomers as expected and similar intensities of α-syn were detected in the α-syn only and α-syn aggregated with Aβ42 fibrils sample.

### α-syn aggregation in α-syn–Aβ42 co-incubation occurs via heterogeneous primary nucleation

The data discussed so far indicate that Aβ42 oligomers trigger the aggregation of α-syn. To further investigate this aggregation mechanism, we performed ThT fluorescence experiments on solutions containing increasing initial concentrations of monomeric α-syn (ranging from 20 to 140 μM) and 2 μM Aβ42 (**Fig. 4a**).

**Figure 4.**
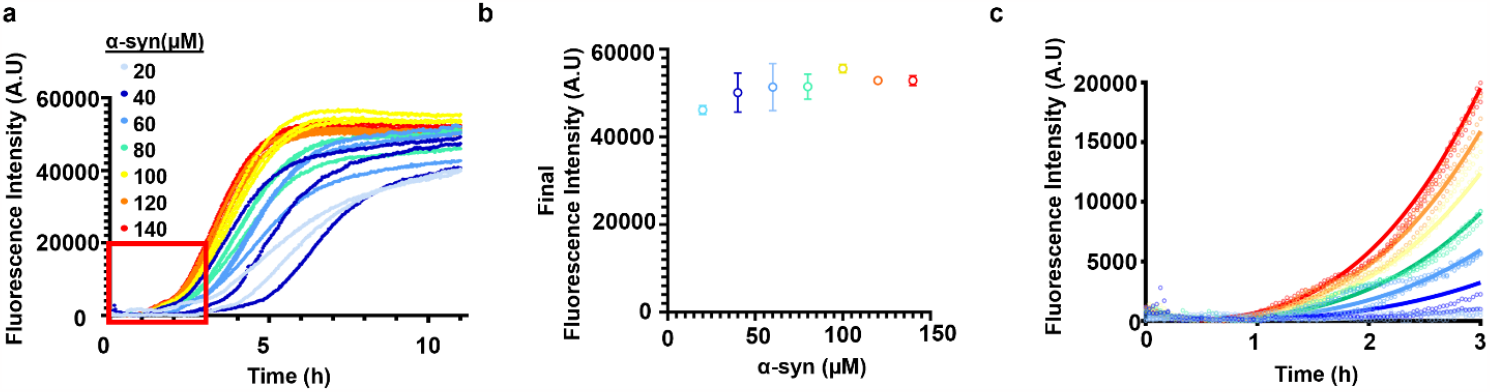
ThT fluorescence measurement of α-syn aggregation in α-syn–Aβ42 co-incubation. (a) ThT fluorescence kinetics of 20–140 μM α-syn and 2 μM Aβ42. The first 3 h were used for global fitting, indicated by the red box. (b) Final ThT fluorescence intensities at 20–140 μM α-syn and 2 μM Aβ42. (c) Global fitting of model parameters for α-syn aggregation. Model parameters C, (n+o_2_) and KM were fitted globally on experimental ThT fluorescence data based on eq. (8). The fitted parameters are C = 2.27·10^10^, (n+o_2_) = 0.69 and K_M_ = 230 μM.

We found that the final ThT fluorescence intensities in the plateau phase do not increase proportionally with the initial α-syn concentrations (**Fig. 4b**), illustrating that not all monomeric α-syn is converted into fibrils, as observed in our immunoblotting analysis (**Fig. 2a, b)**.

In the presence of secondary processes, new nuclei could also form via fibril-dependent nucleation or fragmentation in addition to primary nucleation. In this case the total amount of fibrils is expected to increase linearly with the initial concentration of α-syn (46) as the formation of nuclei and hence monomer depletion until complete consumption is not limited during aggregation. However, the constant final amount of fibrils formed at increasing initial monomeric α-syn suggests that most likely, secondary nucleation processes do not play a substantial role in α-syn aggregation under these conditions.

The lack of secondary processes is further supported by the early-time behavior of the kinetic curves: if secondary nucleation or fragmentation were dominant during aggregation, an exponential increase in the amount of fibrillar α-syn would be expected (48). However, the ThT fluorescence changes polynomially in time (**Fig. S6**) which indicates that most likely primary nucleation is the main source of nuclei (49).

Additionally, we performed Native-PAGE analysis and western blotting on the total protein fraction of the co-incubation samples at the end of aggregation. We found that Aβ42 remained as high molecular weight soluble species that were able to enter the gel in the presence of each of these α-syn concentrations (**Fig. S7**). This result, combined with the observation that neither monomers or fibrils of Aβ42 induce α-syn aggregation, strongly suggests that heterogeneous rather than homogeneous primary nucleation is responsible for the generation of new nuclei. Aβ42 is known to be converted into low-molecular-weight oligomers (50), moreover, it was also reported that α-syn can undergo co-oligomerization with Aβ42 to form hetero-oligomeric species (43). In our case, these oligomers may provide the surface that triggers the aggregation of monomeric α-syn.

To validate our hypothesis, we applied kinetic modelling to describe heterogeneous primary nucleation of α-syn on the surface of the oligomeric species that are generated during the early times of aggregation. Following the approach of Galvagnion et al (46). We limit our analysis to the first 3 hours of aggregation where α-syn and Aβ42 monomer concentrations can be approximated as constant, which facilitates analytical description of aggregation. We describe the generation of oligomers and amyloid aggregation subsequent to α-syn binding to the oligomer surface with differential rate laws. For more details of the mathematical model see Materials and Methods.

Global fitting of the model parameters on eq. (8) (**Fig. 4c**) shows that our kinetic model is consistent with the evolution of the fluorescence signal at the early times of aggregation. We find a weak dependence of oligomer formation and nucleation on monomeric α-syn with an apparent reaction order of 0.69 (o_2_+n in eq. (8) in Materials and Methods) which supports our hypothesis on the heterogeneous nature of primary nucleation (51). We observed saturation of elongation with a Michaelis–Menten-like constant of K_M_ = 230 μM, which agrees well with previously reported K_M_ values of 46–380 μM for lipid vesicles (46,52).

## DISCUSSION

Understanding protein co-aggregation is crucial for developing diagnostic and therapeutic strategies for neurodegenerative diseases. Recently, the identification of LBs in up to 50% of AD patients and Aβ plaques in up to 50% of PD patients has led to research interest on the heterogeneous aggregation of these two peptides.

Although it has previously been reported that α-syn aggregation is triggered by Aβ42 (42) and that Aβ42 aggregation is inhibited by α-syn (39), our findings unify the mechanism of this co-aggregation which has previously remained elusive (**Fig. 5**). Here, we report that when Aβ42 and α-syn are co-incubated *in vitro*, homogeneous amyloid fibrils of α-syn are formed. We also show that under these conditions, Aβ42 is associated to but not part of the core of α-syn fibrils and mainly in the oligomeric conformation. Additionally, we observed that, under these conditions, α-syn aggregates via heterogeneous primary nucleation. Of note, Aβ42 and α-syn hetero-oligomers have also been previously identified (43-45). As fibrils composed of both proteins have not been identified, based on our data we speculate that Aβ42 oligomers and α-syn interact to form hetero-oligomers which likely serve as nucleation sites for the formation of homogeneous α-syn fibrils. The mechanism observed in this work shares remarkable features of α-syn aggregation in the presence of lipid membranes (46) which has also shown to occur via heterogeneous primary nucleation, suggesting that this could be a global property of α-syn when there is a suitable surface to trigger its aggregation. The physicochemical parameters of these suitable surfaces remain unclear, although hydrophobicity is likely to play an important role, considering also that air/water interface is a well-known trigger of α-syn aggregation (53).

**Figure 5.**
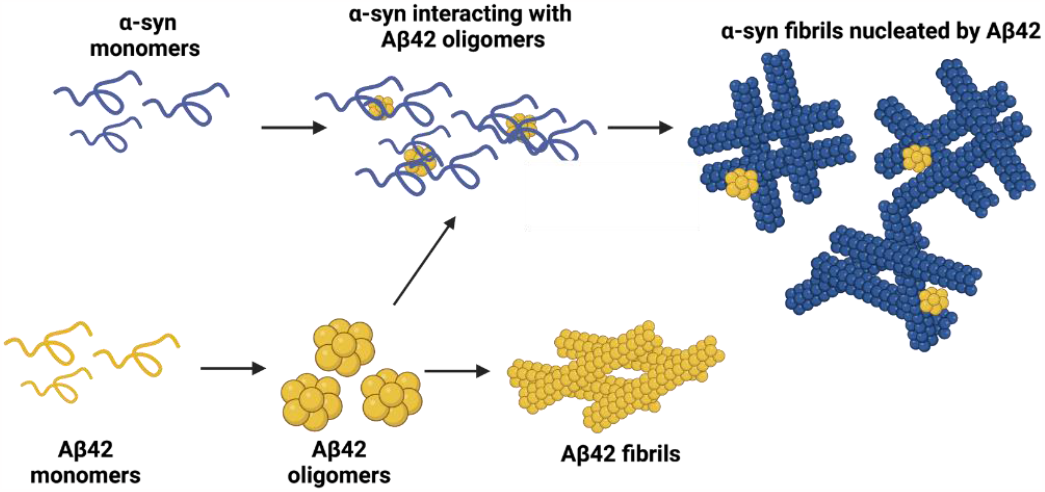
Proposed model for the co-aggregation of Aβ42 and α-syn. When in isolation, Aβ42 undergoes a complete aggregation process that leads to the formation of mature amyloid fibrils. In the presence of α-syn, Aβ42 fibril formation is inhibited. We speculate that, once Aβ42 oligomers are formed, these can interact with α-syn monomers. These heterogeneous oligomers serve as nucleation sites for α-syn aggregation and catalyze the formation homogeneous α-syn fibrils via primary nucleation. Schematic created on Biorender.

It is important to note that α-syn in bulk is not highly aggregation-prone. In fact, in general, the protein requires the interaction with other biomolecules to aggregate (46,54-57). Therefore, Aβ42 oligomers triggering α-syn aggregation is not surprising and a similar stoichiometric relationship has been reported for α-syn and exosomes (58) and glycosaminoglycans (59).

It is plausible that the interaction of α-syn and Aβ42 drives Aβ42 to form highly stable, off-pathway, oligomers that could have additional toxic effects besides triggering α-syn aggregation (60). We believe triggering the aggregation of α-syn could be regarded as an additional toxic mechanism of Aβ42 oligomers. While oligomers of Aβ are widely accepted as the neurotoxic species in AD, we show that it is important not to overlook their consequences in other disease contexts.

## MATERIALS AND METHODS

### Expression and purification of Aβ42

Purification of Aβ42 was carried out as previously described (47). Briefly, the Aβ42 peptide conjugated with the spider silk domain (known as the fusion protein, 20 kDa) was expressed by heat-shock transformation in BL21 *Escherichia coli* (*E. Coli*). Cells were grown in LB supplemented with kanamycin (50 μg/ml) at 37°C with shaking at 200 rpm until they reached an OD of 0.8 and were then induced overnight with 0.1 mM IPTG at 20°C with shaking at 200 rpm. Cells were collected the following day by centrifugation and the pellet was resuspended in 20 mM Tris-HCl, 8 M Urea, pH 8. The resuspended cells were sonicated on ice for 20 minutes (15s on and 45s off pulses, 20% amplitude) and centrifuged once more to clear cellular debris. The supernatant was filtered using a 0.22 μm filter and loaded onto two HisTrap HP 5 ml columns (Cytiva, Little Chalfont, UK) in tandem that had been pre-equilibrated with 20 mM Tris-HCl, 8 M Urea, pH 8 supplemented with 15 mM imidazole (binding buffer). Following sample application, the columns were washed with several column volumes of binding buffer. The fusion protein was then eluted with 5 column volumes of 20 mM Tris-HCl, 8 M Urea, pH 8 supplemented with 300 mM imidazole (elution buffer). This was then collected and dialyzed overnight against 20 mM Tris-HCl, pH 8. After dialysis, the concentration of the fusion protein was measured using a nanodrop. TEV protease was added to the fusion protein at a 1:15 molar ratio overnight at 4°C. Following this, 7 M Guanidine-HCl was added to the sample and incubated on ice for 2 h before applying the sample on to a Superdex 75 Increase pg 10/600 column (Cytiva, Little Chalfont, UK) pre-equilibrated with 20 mM phosphate buffer supplemented with 200 μM EDTA, pH 8 for size-exclusion chromatography. Peaks were collected manually. The concentration of monomeric Aβ42 was determined from the chromatogram using the following calculation:

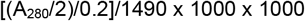

Where 0.2 is the pathlength (cm) of the ATKA Pure (Cytiva, UK), 1490 M^-1^cm^-1^ is the molecular co-efficient of Aβ42.

The stock concentration was diluted to 2 μM for experiments in 20 mM Phosphate buffer supplemented with 200 μM EDTA, pH 8. For Aβ42 fibrils, monomeric Aβ42 was incubated for 24 h after size-exclusion at 37°C.The fibrils were collected by centrifugation at max speed for 30 min after which the concentration of the supernatant was measured with a Nanodrop (Thermo Fisher Scientific, Waltham, MA, USA). The concentration of the supernatant was subtracted from the initial monomeric Aβ42 concentration to obtain the fibril concentration. These were then diluted to 2 μM for experiments in 20 mM Phosphate buffer supplemented with 200 μM EDTA.

### Expression and purification of α-syn

The pT7-7 α-syn WT plasmid (a gift from Hilal Lashuel, Addgene, Watertown, NY, United States (61)), with codon Y136 mutated from TAT to TAC to prevent cysteine misincorporation, was transformed into BL21-Gold (DE3) competent E. coli (Agilent Technologies, Santa Clara, CA, USA) according to the manufacturer’s instructions. Expression of α-syn was induced using 1 mM IPTG at 28°C overnight, ampicillin (100 μg/ml) was included as necessary. The cells were harvested by centrifugation and resuspended in buffer A (20 mM Tris-HCl, 1 mM EDTA, pH 8.0) and protease inhibitors (Roche, Basel, Switzerland). α-Syn was then purified as previously described (62).

### ThT fluorescence assays

20-100 μM α-syn, 2 μM freshly purified monomeric or fibrillar Aβ42 and the two peptides incubated together were prepared with a final concentration of 10 μM ThT dye, gently vortexed and pipetted into non-binding surface black 96-well plates (Greiner Bio-One, Frickenhausen, Austria) in triplicates. The plate was read in a ClarioStar Plus microplate reader (BMG LabTech, Ortenberg, Germany) at 37°C. The excitation and emission wavelength were set to 440nm and 480nm respectively and fluorescent intensity measurements were taken using spiral averaging (3 mm diameter). Buffer only values were subtracted from the sample readings. Readings were taken every 2-5 min. The data were plotted using GraphPad Prism version 9.3.1 for Windows (GraphPad Software, San Diego, CA, United States)

### Immunogold labelling and negative stain transmission electron microscopy

Samples were prepared and incubated for 24 h at 37°C before 4 μl were spotted onto a Formvar/Carbon coated 300 mesh copper grids for 1 minute. Excess sample was removed by blotting dry with Whatman filter paper and the grid was allowed to dry to 2 min. Samples for negative stain TEM were then washed with 4 μl water and stained with 4μl 2% w/v uranyl acetate. Samples for immunogold labelling were blocked using normal goat serum—1:10 dilution in PBS+ [1% BSA, 500 ml/L Tween-20, 10 mM Na EDTA, and 0.2 g/L NaN3] for 15 min after which the grids were incubated on a 20 μl drop of primary antibodies—1:10 dilution of 6E10 (Biolegend, San Diego, CA, USA) and anti-αSyn MJFR1 (Abcam, Cambridge, UK) in PBS at room temperature for 2 h. The grids were then washed with PBS+ three times for 2 min and incubated on a 20 ul drop of secondary antibodies conjugated with gold particles (1:20 dilution of anti-mouse and anti-rabbit secondary antibodies conjugated with 10nm and 6nm gold particles respectively, Abcam, Cambridge, UK). Finally, the grids were washed fives with PBS+ and five times with water for 2 min before being stained with 2% w/v uranyl acetate. Grids were imaged on a T12 Spirit electron microscope (Thermo Fisher Scientific (FEI), Hillsboro, OR, USA). Fibril width was measured using Fiji. All data were plotted using GraphPad Prism version 9.3.1 for Windows (GraphPad Software, San Diego, CA, United States). 6nm and 10nm gold particles were assigned colors (red and blue, respectively) after imaging using an in-house Python script using the morphology module of SKimage.

### Dot blotting

Dot blots were carried out on samples that were aggregated in the microplates without ThT at the endpoint of aggregation. Samples were collected and centrifuged at max speed (∼17,000 g) for 30 min on a benchtop centrifuge to separate the soluble and insoluble aggregates. 3-5 repeats of each sample were spotted onto 0.45 μM nitrocellulose membrane and blocked in 5% non-fat milk in 0.1% PBS-Tween for 1 h at RT. The membranes were then incubated in primary antibodies (1:1000 dilution in 0.1% PBS-Tween for both anti-α-syn and 6E10) overnight at 4°C under constant shaking. The following day, the membranes were washed three times for 10 min each in 0.1% PBS-Tween. Membranes were then incubated in secondary antibodies conjugated with an AlexaFluor tag (anti-mouse 647 and anti-rabbit 555, diluted 1:2000 and 1:5000 0.1% PBS-Tween, respectively, Thermo Fisher Scientific, Waltham, MA, USA) at room temperature for 1 hour, protected against light. Following three further washes for 10 min each in 0.1% PBS-Tween, the membranes were detected with the appropriate laser using the Typhoon scanner (GE Healthcare, Amersham, UK).

### PK digestion

Fibrils were collected from the co-incubation sample at the end of aggregation by centrifugation at max speed (∼16,000g) for 1h. α-Syn fibrils were collected after two weeks of incubation at 37°C under quiescent conditions. Fibrils were resuspended in buffer and treated with 0-15μg/ml of PK for 20mins. Sample were then incubated at 95°C for 5 minutes to stop the enzymatic reaction and samples were prepared for SDS-PAGE and western blotting analysis. The primary antibodies used for this analysis are detailed in Table S1 and were purchased as a kit from Cosmo Bio Co., LTD (Japan, Tokyo). Densitometry analysis was carried out using the Fiji software and the IC50 was calculated from normalized data using the GraphPad Prism (GraphPad Software, CA, USA).

### SDS-PAGE and western blotting

Fibrils from the co-incubation sample were collected by centrifugation at max speed for 30 min and the supernatant (S1) was removed and separated from the pellet (P1). The pellet was washed with 50 μl 2% SDS and centrifuged. The supernatant was again removed (S2) and the pellet was resuspended in 50 μl buffer to remove any residual SDS (P2), the pellet was centrifuged, and the supernatant (S3) and pellet were separated (P3) after which the pellet was treated with 50 μl 4 M guanidine hydrochloric acid (G-HCl) for 2 h at RT before SDS-PAGE and western blot analysis. Samples were prepared in 4X LDS sample buffer and 10X reducing agent after which they were boiled at 95°C for 5 min. Samples were then run on 4-12% Bis-Tris NuPAGE gels (Thermo Fisher Scientific, Waltham, MA, USA) and transferred onto a 0.45 μm nitrocellulose membrane for 7 min at 20 V with the i-Blot 2 (Thermo Fisher, Waltham, MA, USA). Blocking, incubation with antibodies and detections were carried out as described above.

### Native-PAGE and western blotting

Samples were prepared and aggregated in 96-well microplates with a non-binding surface for 24 h after which 20 μl of each sample was prepared in native sample buffer and run on Novex Tris-Glycine gels (Thermo Fisher Scientific, Waltham, MA, USA) in native running buffer as per the manufacturer’s instructions. The gel was then transferred onto a 0.45 μM nitrocellulose membrane using an iBlot2 (Thermo Fisher Scientific, Waltham, MA, USA) for 7 min at 25 V. The membrane was then blocked, incubated with primary and secondary antibodies, and imaged as described above.

### Modelling of aggregation kinetics

#### Kinetic model of aggregation

We consider the generation of oligomers which can be composed of both Aβ42 and α-syn. α-syn monomers then bind to the surface of oligomers where they form fibril-competent nuclei. We derive differential rate equations describing the number concentration of oligomers *O* (eq. (1)) and α-syn fibrils *F* (eq. (3)) as well as the fibril mass concentration *M* (eq. (4)). The rate of oligomer formation can be described by eq. (1):

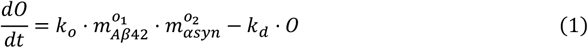

where *m*_*Aβ*42_ and *m*_*αsyn*_ are the concentration of monomeric Aβ42 and α-syn, *o*_1_ and *o*_2_ are the apparent reaction order of oligomer formation relative to monomer concentration of Aβ42 and α-syn, respectively, *k*_*o*_ and *k*_*d*_ denote the rate constant of oligomer formation and dissociation. We note that considering homo-oligomerization of Aβ42 is a special case of eq.

(1) where *o*_2_ = 0. Although in principle the dissociation of oligomers may be a function of their surface coverage *Φ*, the large excess of a-syn even in the lowest α-syn-Aβ42 ratio suggests that the coverage remains close to 1 at all α-syn concentrations. Hence, *k*_*d*_ is assumed constant in our kinetic model.

In the early time limit *m*_*Aβ*42_ and *m*_*αsyn*_ is approximated by their initial concentrations, and hence eq. (1) can be solved analytically for *O*:

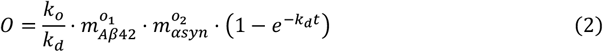

We hypothesize that binding of monomeric α-syn happens on a much shorter time scale then the other processes, hence binding equilibrium is reached instantaneously. Moreover, due to the large excess of α-syn compared to Aβ42 saturated surface coverage is assumed on the oligomers, that is, Φ ≈ 1 in all cases. Subsequent to binding, heterogeneous primary nucleation occurs on the surface of oligomers. The rate of nuclei formation is written as:

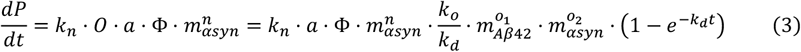

*a* denotes the number of binding sites of α-syn on the surface of one oligomer, *n* is the apparent reaction order of nucleation relative to monomeric α-syn. Since secondary processes were assumed to be negligible at these conditions, substituting eq. (2) into eq. (3), and subsequent integration yields:

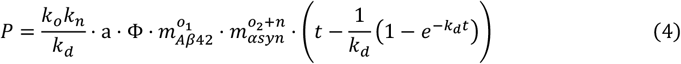

The fibril mass increases over time via elongation:

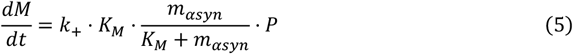

Integration of eq. (5) provides the total fibril mass at time point *t*:

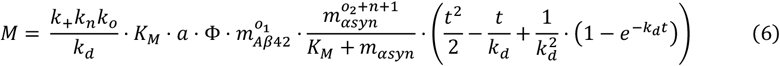

Which can be further simplified upon expansion of the exponential expression into:

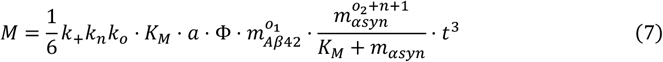

Assuming linear dependence of the measured fluorescence intensity on the fibril mass concentration (*F*.*I*. = *λ* ⋅ *M*) we get:

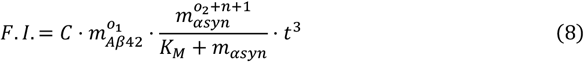

Where *C*, (*o*_2_ + *n* + 1) and *K*_*M*_ were evaluated by a global fit to the experimental ThT profiles curves by MATLAB homemade programs.

## Data Availability

All data are contained within the main text and supplementary information.

## Supporting Information

This article contains supporting information.

## Acknowledgments

We thank Dr Henrik Biverstål (Karolinska Institutet) for providing the expression plasmid for the silk domain-Aβ42 fusion protein and the Electron Microscopy Centre facilities at The Centre of Structural Biology for TEM experiments. We thank Prof. Louise Serpell (University of Sussex) for helpful discussions.

## Author Contributions

D.M.V. and F.A.A. designed the research. D.M.V, R.J.T and Y.J performed the experiments. D.M.V and F.A.A. wrote the manuscript. M.P. and P.A. performed the kinetic analysis. All authors analyzed the data and commented on the manuscript.

## Funding and additional information

We thank UK Research and Innovation (Future Leaders Fellowship MR/S033947/1), the Engineering and Physical Sciences Research Council (grant EP/S023518/1), the Alzheimer’s Society, UK (Grant 511) and Alzheimer’s Research UK (ARUK-PG2019B-020) for support. RJT was supported by a scholarship by the Department of Chemistry (Imperial College London). P.A. kindly acknowledges the European Research Council through the Horizon 2020 research and innovation programme (grant agreement No. 101002094) for financial support.

## Conflict of Interest Statement

The authors declare that they have no conflicts of interest with the contents of this article.

## Open access Statement

For the purpose of Open Access, the author has applied a CC BY public copyright license to any Author Accepted Manuscript version arising from this submission.

## Notes

### Competing Interest Statement

The authors have declared no competing interest.

### Summary of Updates

Formatting and changes to text

