## Supplementary Data for "α-Synuclein Aggregation is Triggered by Amyloid-β Oligomers via Heterogeneous Primary Nucleation"

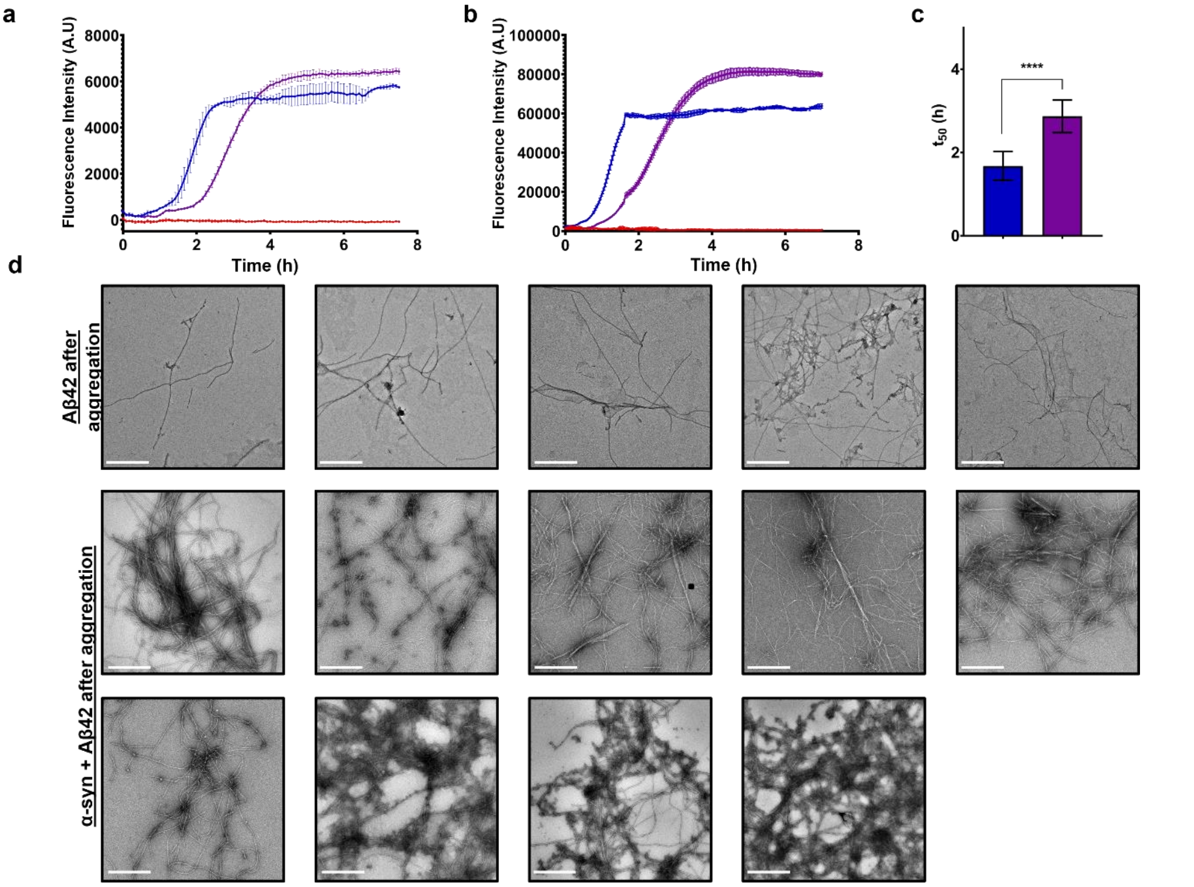


**Fig. S1. Aggregation Aβ42 and α-syn aggregated with Aβ42**. (a-b) Two independent aggregation assays. α-syn is shown in red, Aβ42 is shown in blue, and α-syn incubated with Aβ42 is shown in purple (c) The t50 of 3 independent aggregation repeats were averaged. Error bars are shown as SD. Unpaired, non-parametric Mann Whitey's test, where p = 0.1234 (ns), 0.0332 (*), 0.0021 (**), 0.0002 (***) and <0.0001 (****). (d) Negative stain TEM. All TEM images used to confirm fibril formation of Aβ42 and α-syn incubated with Aβ42 at the end of aggregation. Scale bar representative of 0.2µm.

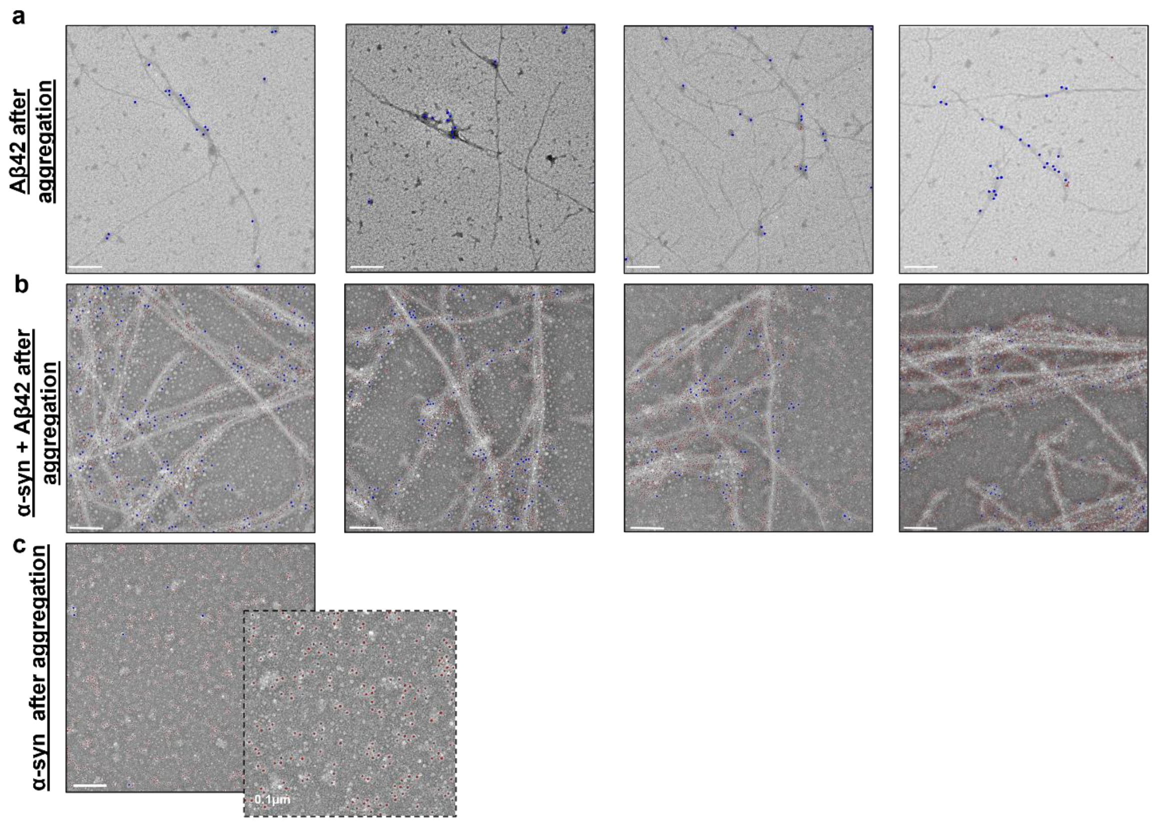


**Fig. S2. Immunogold labelling and negative stain TEM.** All images used to quantity fibril length of (a) Aβ42 and (b) the co-incubation samples after aggregation. Scale bars representative of 0.2μm unless stated otherwise. (c) α-syn after aggregation is also shown. Red 6nm gold particles bind to the anti-α-syn primary antibody, and blue 10nm gold particles bind to the anti-Aβ42 6E10 primary antibody.

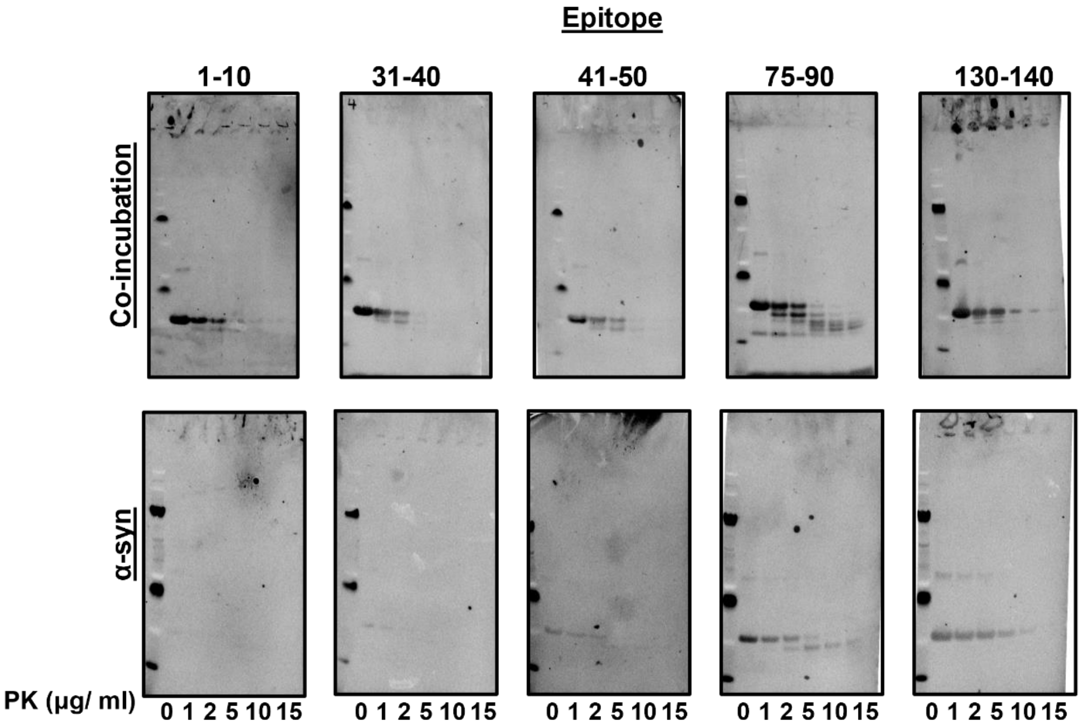


**Fig. S3. Proteinase K digestion of α-syn fibrils in the presence and absence of Aβ42.** Fibrils were collected at the end of aggregation and treated with increasing concentrations of proteinase K for 20 minutes. SDS-PAGE and western blotting with antibodies scanning the sequence of α-syn reveal a similar PK digestion profile for fibrils formed in both conditions.

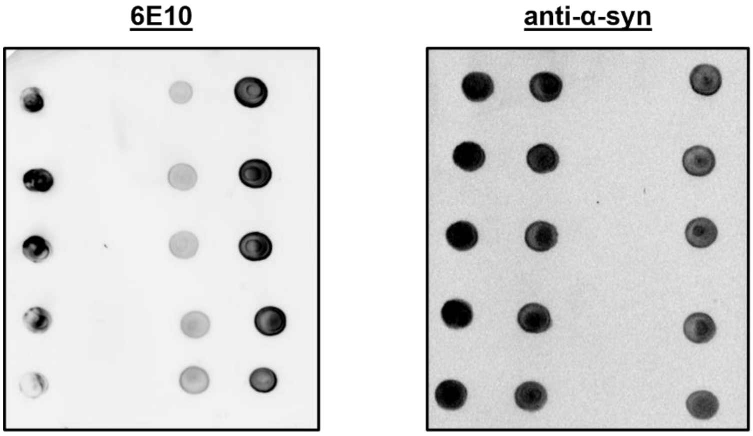


**Fig. S4. Solubility of Aβ42 and α-syn after co-incubation.** Dot blot analysis on the soluble fractions of aggregated samples detected with 6E10 (left) and anti-α-syn (right) primary antibodies. 5 repeats for each sample were quantified (**Fig. 2b**)

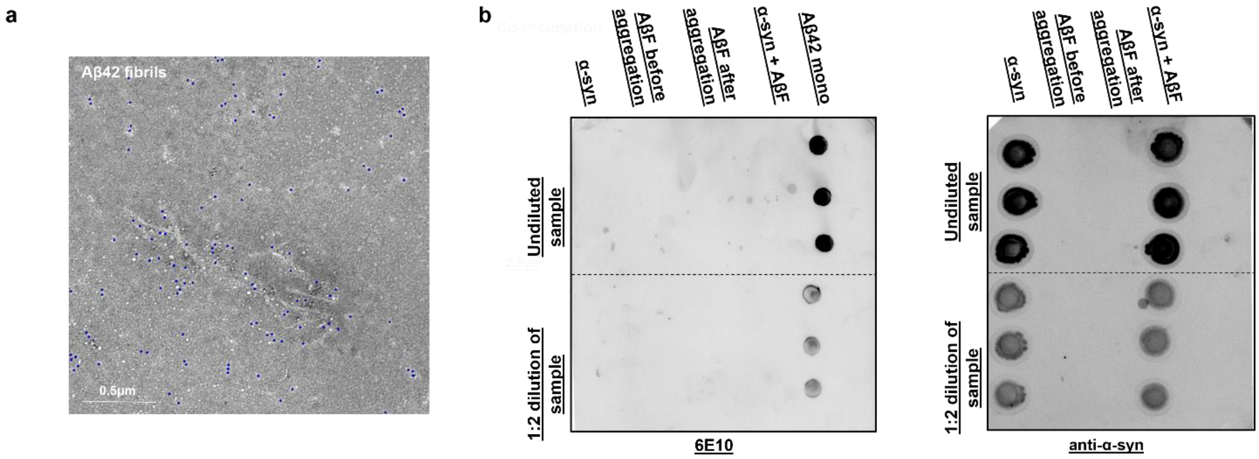


**Fig. S5. AβF do not nucleate the aggregation of α-syn.** (a) Immunogold labelling Aβ fibrils alone at the end point of aggregation. Fibrils highly decorated with 10nm gold particles (Ab–6AuNPs, red) specific for Aβ42 as opposed to 6nm gold particles labelling α-syn (Ab–10AuNPs, blue). (b) Uncropped dot blot analysis shown in Fig. 3d. Dot blot analysis was carried out on the soluble fractions of aggregated samples detected with 6E10 (left) and anti-α-syn (right) primary antibodies. No soluble Aβ42 was detected in any sample except freshly purified Aβ42 monomers as expected and similar intensities of α-syn were detected in the α-syn only and α-syn aggregated with Aβ42 fibrils sample.

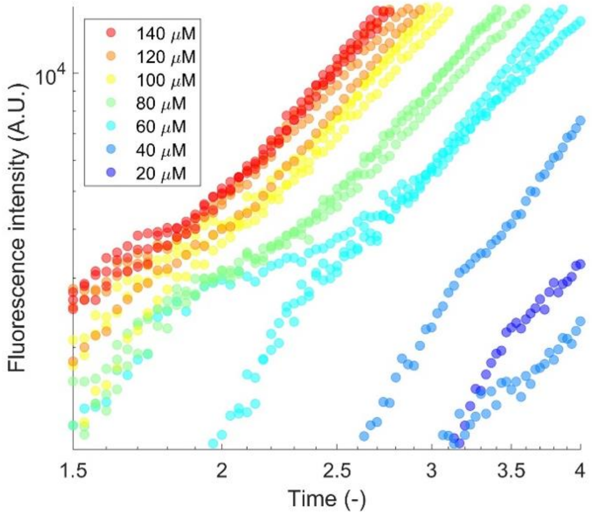


**Fig. S6. ThT fluorescence kinetics of the aggregation of α-syn.** The fluorescence signal increases linearly on the logarithmic scale plot implying polynomial time evolution of the total fibril mass in the first 4 hours of co-incubation of Aβ42 and α-syn.

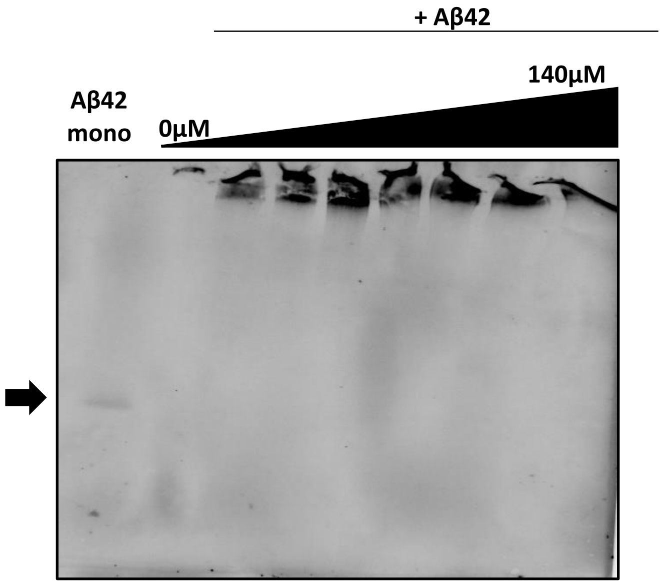


**Fig. S7 Aβ42 remains soluble at increasing molar ratios when incubated with α-syn.** Native-PAGE and western blot analysis revealed that compared to monomeric Aβ42, Aβ42 after aggregation was detected only has aggregates stuck in the well of the gel. In the presence of increasing α-syn concentrations of 20-100 µM), Aβ42 remains as soluble high molecular weight assemblies.

| Epitope | Sequence |
|---|---|
| **1-10** | MDVFMKGLSKC |
| **31-40** | GKTKEGVLYVC |
| **41-50** | GSKTKEGVVHC |
| **75-90** | CTAVAQKTVEGAGSIAAA |
| **130-140** | CEGYQDYEPEA |

**Table S1.** Epitopes of the anti-α-syn antibodies used in the PK digestion analysis shown in Fig.1e.
